# Differential Sensitivity of Impedance Plethysmography and Photoplethysmography Sensors to Temperature-Induced Peripheral Vasoconstriction

**DOI:** 10.1101/2025.05.27.656391

**Authors:** Seobin Jung, Seamus Thomson, Alexandros Pantelopoulos, Lindsey Sunden, Pete Richards, Shwetak Patel, Sam Sheng

## Abstract

Impedance plethysmography (IPG) and photoplethysmography (PPG) are non-invasive techniques for measuring blood volume changes. This study investigated the differential responses of IPG and PPG to temperature-mediated vasoconstriction induced by localized cooling. Twenty-one participants underwent control and treatment conditions, with real or fake ice cubes applied to the forearm. PPG signal amplitude significantly decreased with cooling (p < 0.001), indicating sensitivity to capillary blood flow changes. In contrast, IPG signal amplitude remained stable, suggesting it primarily reflects blood flow in larger/deeper vessels. Blood pressure remained stable, while heart rate decreased. These findings suggest IPG is less sensitive to capillary-level changes than PPG and may be more suitable for monitoring deeper blood flow. This study provides insights into the distinct sensitivities of IPG and PPG, with implications for wearable device development and cardiovascular monitoring.

## 1 Introduction

IPG is a non-invasive technique for measuring blood flow in the periphery [1]. Initially developed for monitoring vascular conditions such as deep vein thrombosis and peripheral arterial disease [2], IPG has more recently been employed in wearable sensor research [3]. The sensing mechanism of IPG involves applying an alternating current between two outer electrodes, and measuring the resulting voltage potentials with two inner electrodes, producing time-series impedance data (Fig. 1). While traditionally confined to clinical settings for assessing vascular health, advancements in wearable sensor technology have broadened the potential applications of IPG. More recently, some studies report using wearable IPG signal input for the passive monitoring of pulsatile waveforms in the periphery and the development of cuffless blood pressure estimation models [3, 7].

**Figure 1.**
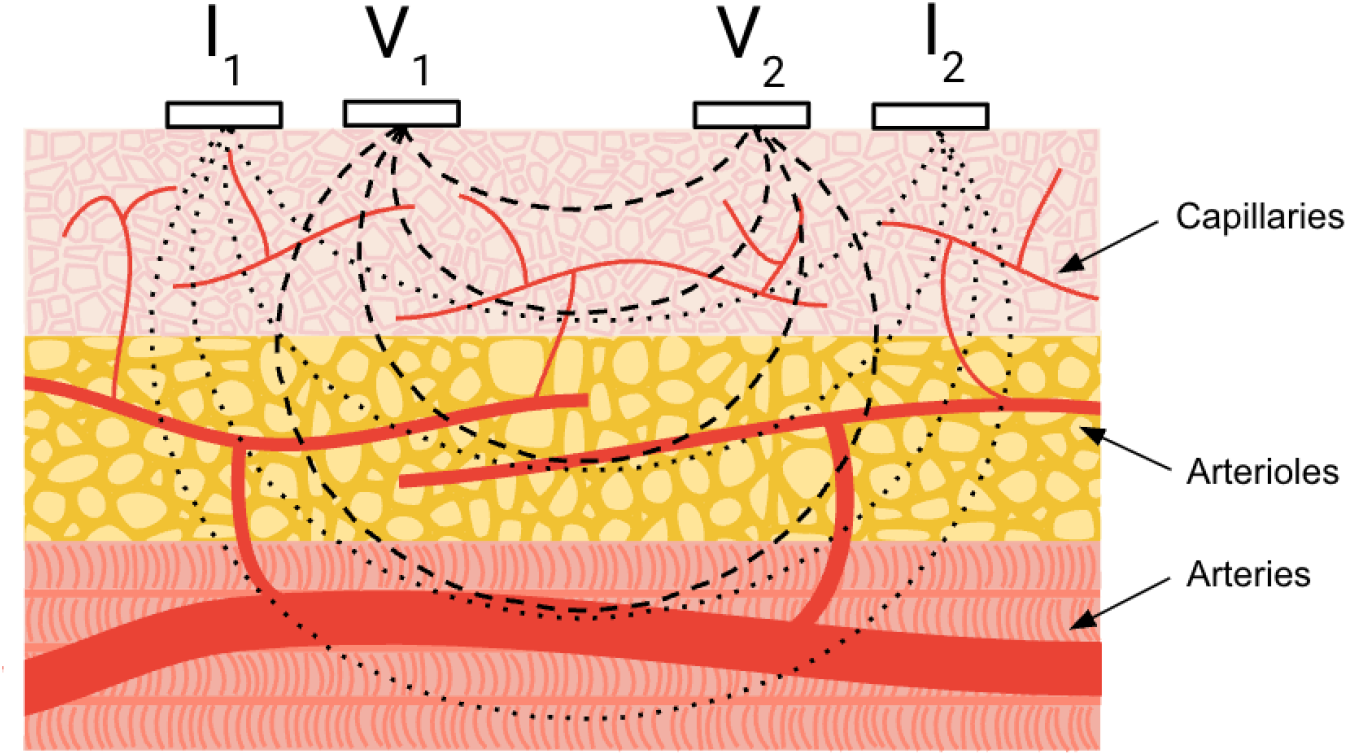
Cross-sectional representation of four-electrode impedance plethysmography setup for measuring blood volume variations. Current injection electrodes (I1 and I2) establish an electric field within the stratified tissues. Voltage sensing electrodes (V1 and V2) measure the resulting impedance, which is modulated by pulsatile blood flow in the underlying vascular structures. Dotted lines approximate the current flow, dashed lines conceptually indicate the region influencing the impedance measurement.

However, a deeper understanding of the sensitivity of IPG to different types of blood flow -particularly in smaller blood vessels such as capillaries - remains an area of ongoing research and crucial for interpreting IPG signals in wearable devices. This understanding is particularly important in the context of vasoconstriction - a physiological response that primarily affects capillary blood flow - and how this impacts IPG signals in wearable devices. Photoplethysmography (PPG), another non-invasive technique which optically measures pulsatile signal, is known to be sensitive to capillary blood flow, with signals measured on the skin surface reflecting blood flow and diffusion predominantly in the capillary beds rather than larger and deeper arteries [4]. In contrast, the extent to which capillary blood flow influences IPG signals remains unclear.

Temperature-mediated vasoconstriction, a well-documented physiological phenomenon, has been extensively-covered in the literature [5, 6]. More specifically, exposure to cold stress causes a reduction in PPG peak amplitudes due to the narrowing of smaller blood vessels. By positioning a PPG sensor between two IPG sensing electrodes on top of the radial artery and inducing a decrease in skin temperature, we can simultaneously observe the effects of vasoconstriction on both IPG and PPG signals.

This study aims to investigate the effects of reduced capillary blood flow on IPG signals, comparing them to PPG signals which are known to be sensitive to capillary blood flow. By inducing localized vasoconstriction through localized cooling, we aim to discern the distinct responses of IPG and PPG signals to changes in capillary blood flow. This investigation will contribute to a more nuanced understanding of IPG and its potential for broader wearable applications.

## 2 Methodology

### 2.1 Participants

Participants were recruited from a pool of Google full-time employees aged 18 years or older, located within the United States of America, capable of giving informed consent, wearing test devices for extended periods, and participating in test protocols while wearing validated research devices. Participants were excluded if they were unwilling or unable to provide informed consent, wear test devices for extended periods of time, unable to participate in test protocols while wearing validated research devices, or have presence of potential identifiers or markings on the wrist region where thermal images are taken.

### 2.2 Instrumentation and Signal Processing

An infrared thermal camera (FLIR E8) served the dual purpose of identifying the radial artery and recording skin temperature. IPG was implemented via gel electrodes (3M 2760-5) positioned with an alignment guide as shown in Fig. 2 right panel, while the PPG sensor was placed between the IPG electrodes. Per participant, the PPG sensor was secured with an elastic band selected for a snug fit that was neither too tight nor too loose to maintain the PPG sensor positioning. An automated blood pressure cuff (Omron BP7450) was placed on the opposite arm, and additional electrodes were applied to form electrocardiography (ECG) lead I configuration (left shoulder, right shoulder, and right leg ankle). Physiological signals were acquired using a Biopac MP36R system with dedicated modules for IPG (Biopac SS31L, 100 kHz excitation frequency, excitation current = 400 µArms, continuously applied), PPG (Biopac SS4LA, 860 nm ± 60 nm infrared, light emitting diode (LED) current drive = 5 mA continuously driven), and ECG (Biopac SS2LB). For the Biopac PPG module, the LED center to the photodiode center spacing was 3.81 mm.

**Figure 2.**
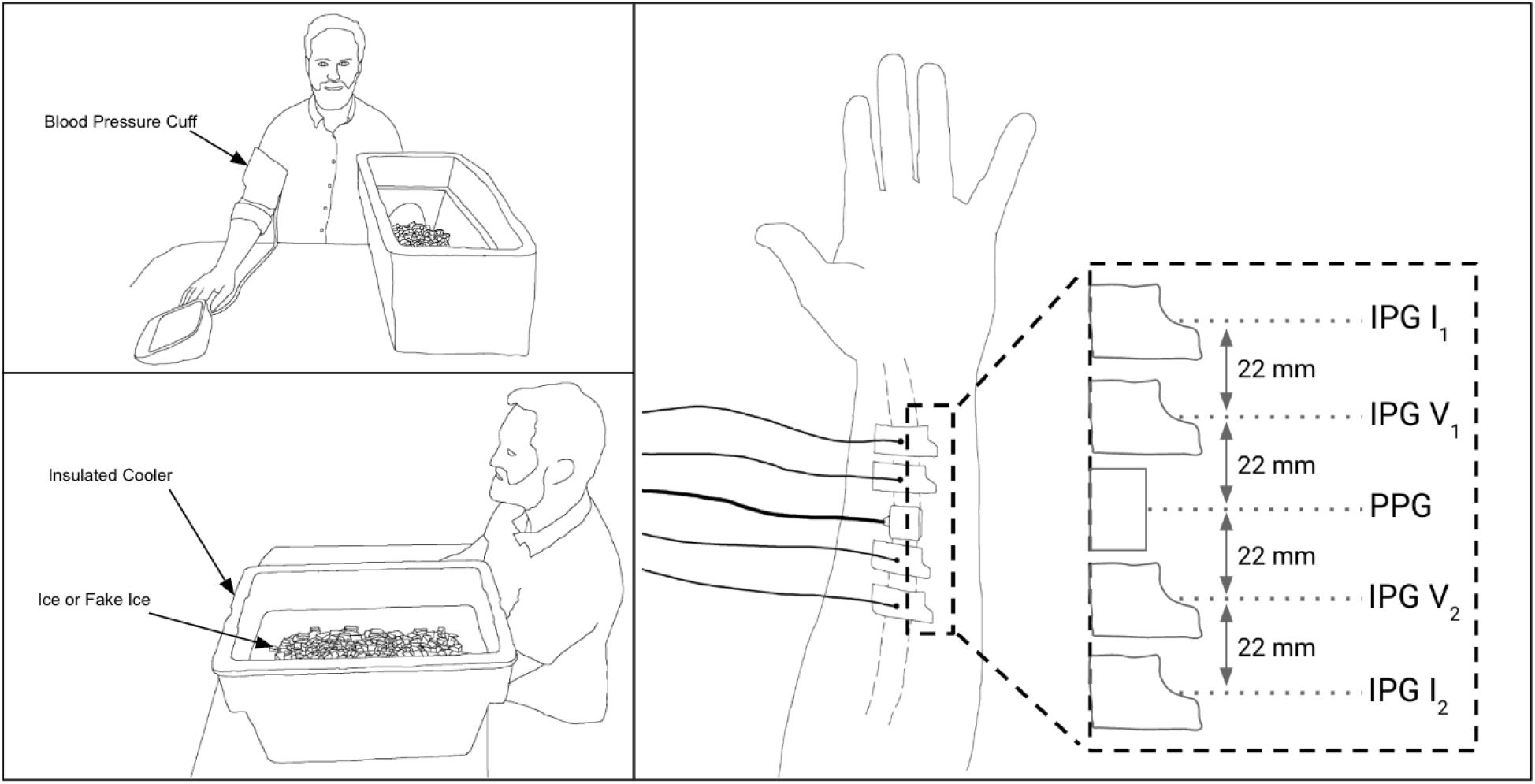
Experimental setup and sensor configurations. The left two panels depict the overall setup including the participant, insulated cooler used for the cold stimulus, and the position for the blood pressure measurement. The right panel shows the placement of four IPG electrodes and PPG module along the forearm targeting the radial artery.

Blood pressure, temperature readings, and sensor data (IPG, PPG, and ECG) were analyzed using Python scripts. The ECG R peaks were found using the Pan-Tompkins algorithm. The Biopac sensor data was recorded at a sampling rate of 2 kHz with time synchronization. Signal processing involved bandpass filtering (0.5 to 12 Hz, Chyebyshev type II, 4th order) for PPG and IPG signals, introducing no phase shift. Since Biopac-reported IPG Z channel was often saturated while its dZ/dt channel was intact (a characteristic attributed to its hardware-based filtering in the IPG module), the IPG signal underwent additional processing. This processing included integration of the dZ/dt channel to derive a Z channel and baseline flattening using a moving average subtraction method with a window size of 2000 samples. Exemplary waveform traces from one participant with the described signal processing are shown in Fig. 3. For statistical testing, paired t-tests were performed using the Python package Scipy.

**Figure 3.**
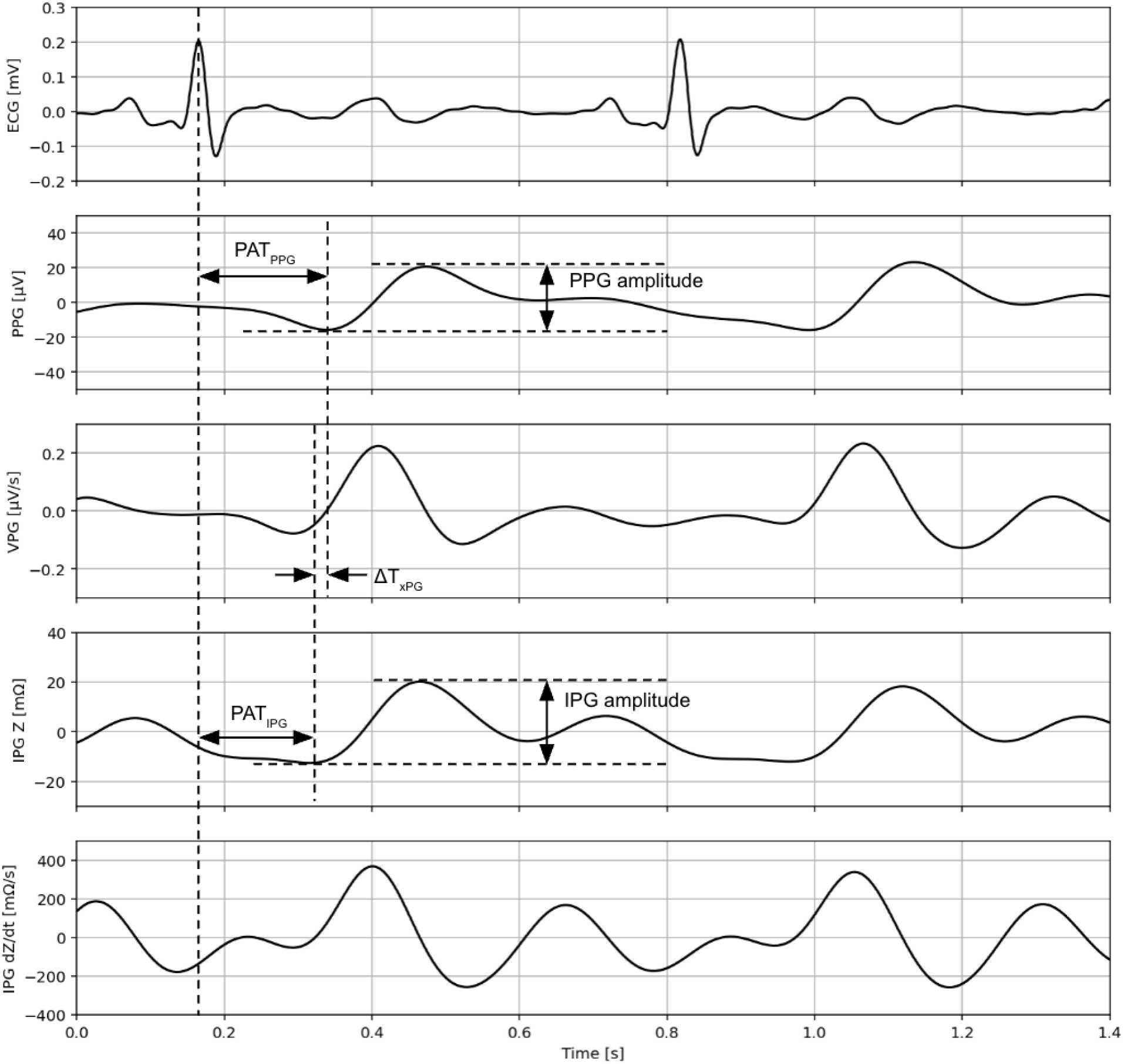
Illustration of feature extraction from simultaneously acquired cardiovascular signals. The traces display ECG, PPG, velocity plethysmography (VPG, first time derivative of PPG), IPG Z, and IPG dZ/dt. Indicated parameters include peak amplitudes for PPG and IPG, and timing features such as pulse arrival time for PPG and IPG referenced to the preceding ECG R peak, and the time interval between PPG valley and IPG Z valley (Δ*T*_*xPG*_).

### 2.3 Procedure

This study employed a within-subject design wherein participants were sequentially exposed to both control and treatment conditions. The procedure was identical for both conditions, with the exception of the stimuli applied. All subjects went through two groups: the treatment procedural group received real ice cubes to induce localized change in temperature, whereas the control procedural group received fake ice cubes to replicate the potential physical disturbances to the sensors such as pressure without the temperature change. Participants were instructed to position their arm with sensors into an insulated cooler on a table, ensuring that the arm remained parallel to the table surface and the ventral wrist facing upwards (Fig. 2 left panel). Throughout the data collection period, participants were instructed to maintain both physical stillness and silence. Following an initial blood pressure measurement and the placement of a towel over the participant’s arm, IPG, PPG, and ECG signals were recorded for a two minute duration. Subsequently, skin temperature was measured at three distinct locations: proximally and distally adjacent to the outer IPG electrodes, and centrally adjacent to the PPG sensor. 1.6L of either real or fake ice cubes, depending on the condition, were then placed on the towel, covering the forearm with particular attention to the sensor area. After four and six minutes, the towel was lifted to retake temperature measurements. Finally, signal acquisition was stopped and a final blood pressure reading was taken. “Baseline” and “Post Intervention” periods were defined as the 60 beats immediately after ice placement and the final 60 beats of signal recording, respectively (Fig. 4).

**Figure 4.**
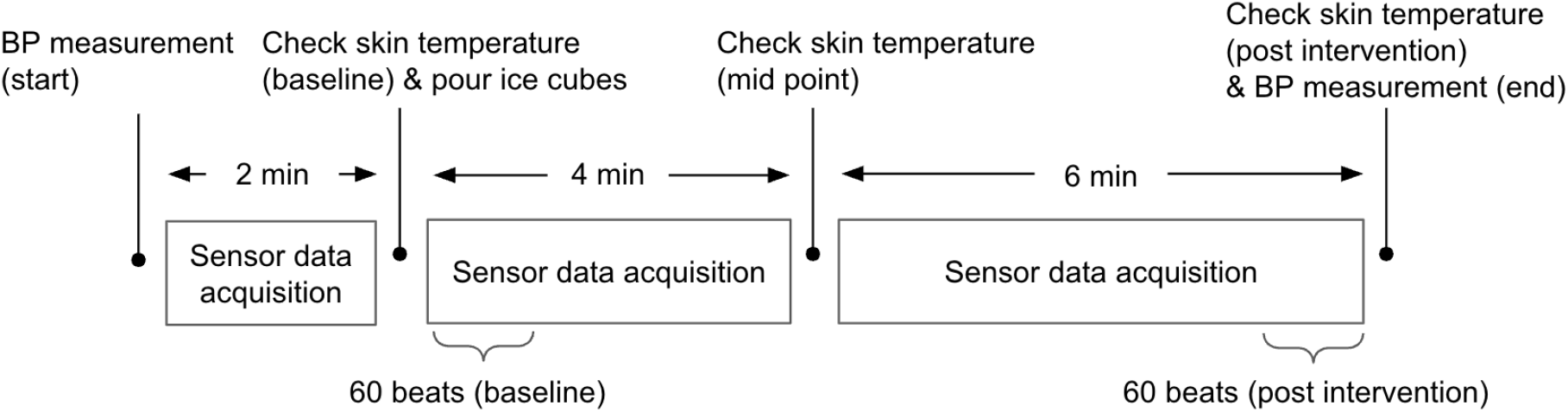
Schematic diagram of the experimental procedure and timeline. Key steps include baseline and post-intervention BP measurements, skin temperature checks, application of cold stimulus, and defined periods of sensor data acquisition. Analysis segments correspond to 60 heartbeats during the start of the baseline and end of the post-intervention phases. The experimental procedure was the same for both the control condition and the treatment condition except the ice cubes being fake vs. real. BP measurements were done at the start of the control and the end of the treatment condition.

### 2.4 Ethical Considerations

All participants provided informed consent prior to their participation in the study. The study protocol was approved by an Institutional Board Review provided by the WIRB-Copernicus Group (WCG) under the protocol number 20243288.

## 3 Results

### 3.1 Participant Demographics

This study included 21 participants, with a mean age of 35.3 ± 6.6 years and a semi-balanced gender distribution (9 females, 12 males). Demographic information is presented in Table 1, including age, gender, height, weight, body mass index (BMI), self-assessed Monk Scale, and self-assessed Fitzpatrick Scale. For continuous values, mean ± standard deviation and range (minimum-maximum) are reported. For categorical values, numbers for each category and its percentage are reported.

**Table 1:** Participant demographics.

| Characteristic | Value |
| --- | --- |
| Age (years) | $35.3 \pm 6.6$ (23-57) |
| Gender (Male/Female) | 12 (57%) / 9 (43%) |
| Weight (kg) | $75.7 \pm 17.4$ (50.0-111.1) |
| Height (cm) | $174.7 \pm 9.2$ (157.5-188.0) |
| BMI ($\text{kg}/\text{m}^2$) | $24.6 \pm 4.2$ (17.2-35.2) |
| Monk Scale | A: 0 (0%), B: 5 (23.8%), C: 2 (9.5%), D: 6 (28.6%), E: 4 (19%), F: 2 (9.5%), G: 2 (9.5%), H: 0 (0%), I: 0 (0%), J: 0 (0%) |
| Fitzpatrick Scale | I: 2 (9.5%), II: 4 (19%), III: 8 (38%), IV: 5 (24%), V: 2 (9.5%), VI: 0 (0%) |

### 3.2 Skin Temperature and Blood Pressure Analysis

The ice cube intervention decreased skin temperature in the treatment group by an average of 13.2 °C across the three measurement locations. In contrast, the control group experienced a slight temperature increase of 0.5 °C, likely due to the insulating effect of the towel (Fig. 5). A paired t-test analysis revealed a statistically significant difference (p < 0.001) in skin temperature between the treatment and control groups (Table. 4, Appendix).

**Figure 5.**
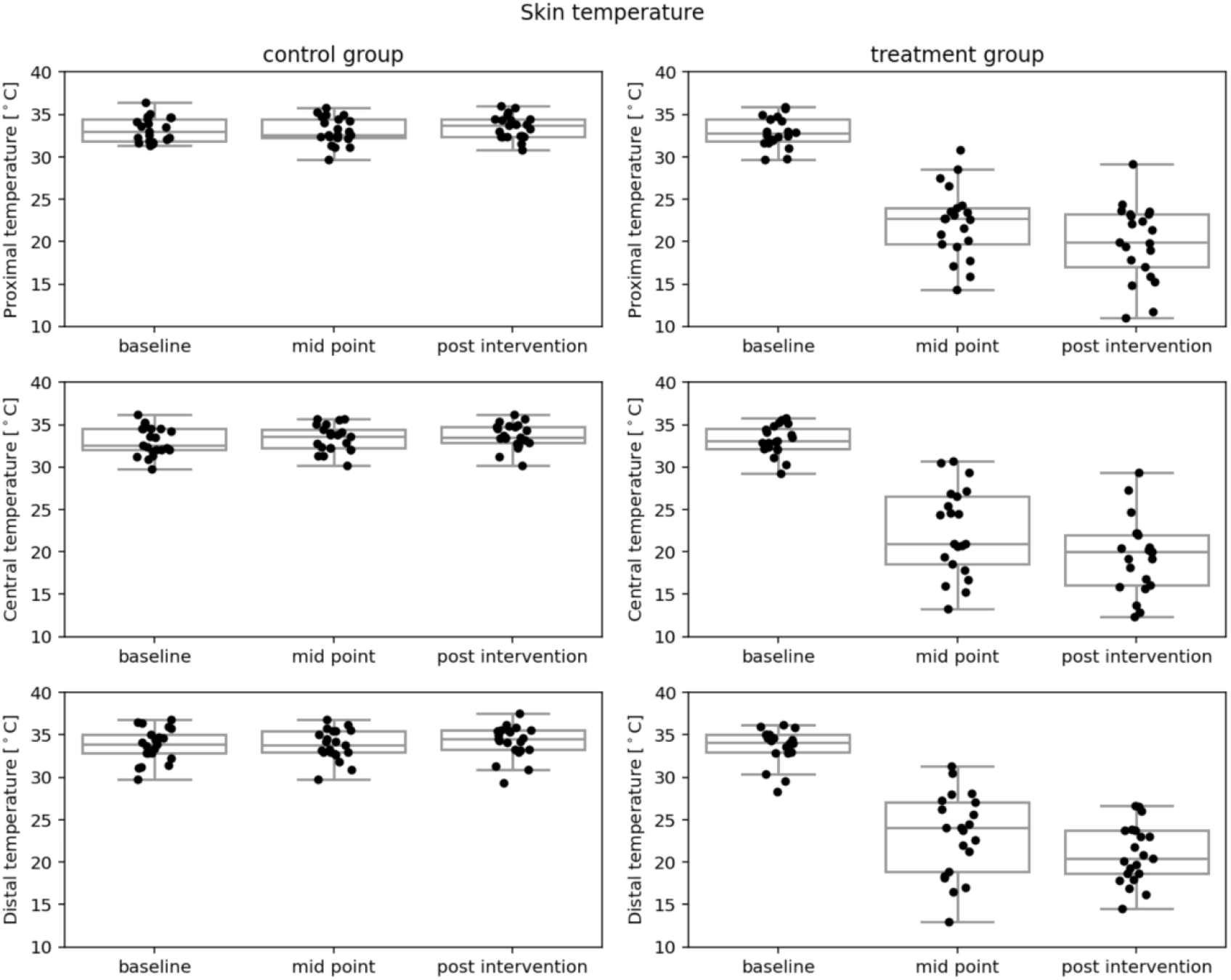
Comparison of proximal, central, and distal skin temperatures [unit: °C] between the control and treatment groups. Measurements were taken at baseline, mid-point of the intervention, and post-intervention. Individual points show data for each participant, box plots indicate the collective distributions.

Blood pressure (systolic, diastolic) did not significantly change between the start and end of the measurement period. Heart rate, however, did significantly decrease by a mean of 2.1 beats per minute (Fig. 6, Table. 5, Appendix).

**Figure 6.**
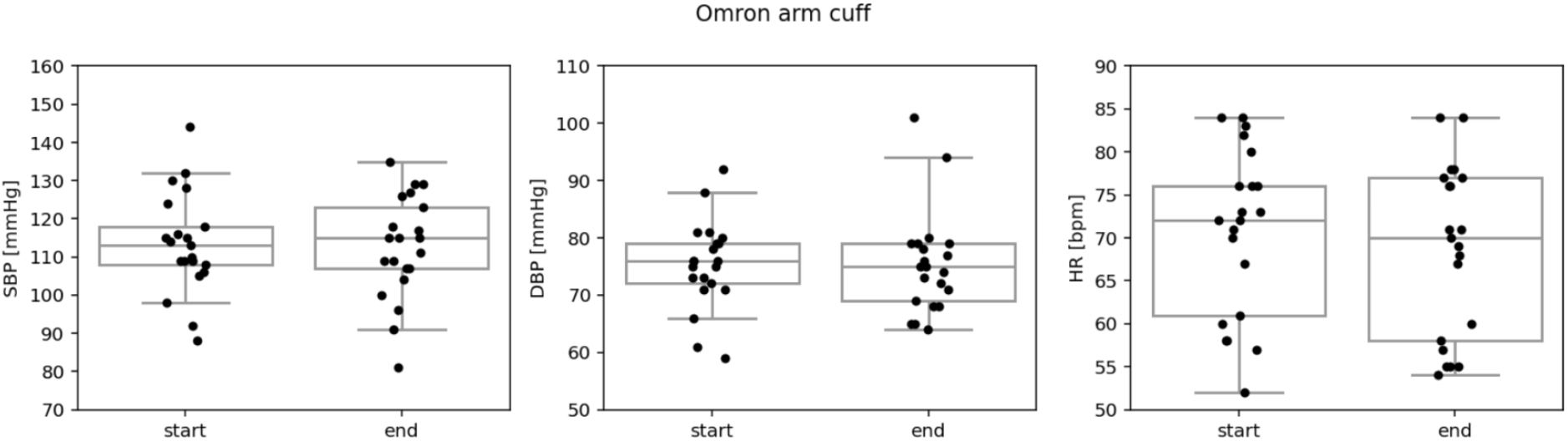
Omron arm cuff measurements of SBP [unit: mmHg], DPB [unit: mmHg], and HR [unit: bpm] at the start and end of the experimental protocol. Individual points show data for each participant, box plots indicate the collective distributions.

### 3.3 Signal Amplitude Analysis

Both PPG and IPG used the same peak finding approach which consisted of being guided by ECG R peaks to scope one beat interval and finding minima (for valley) and maxima (for peak) within the beat interval. PPG and IPG amplitude features using this method are shown in Fig. 3. For each phase of the data collection, peak-to-peak amplitudes were calculated out of 60 beats (Fig. 4). Table 2 reports Mean ± Standard Deviation (SD) of PPG and IPG signal at baseline and post intervention, and across control and treatment groups.

**Table 2:**
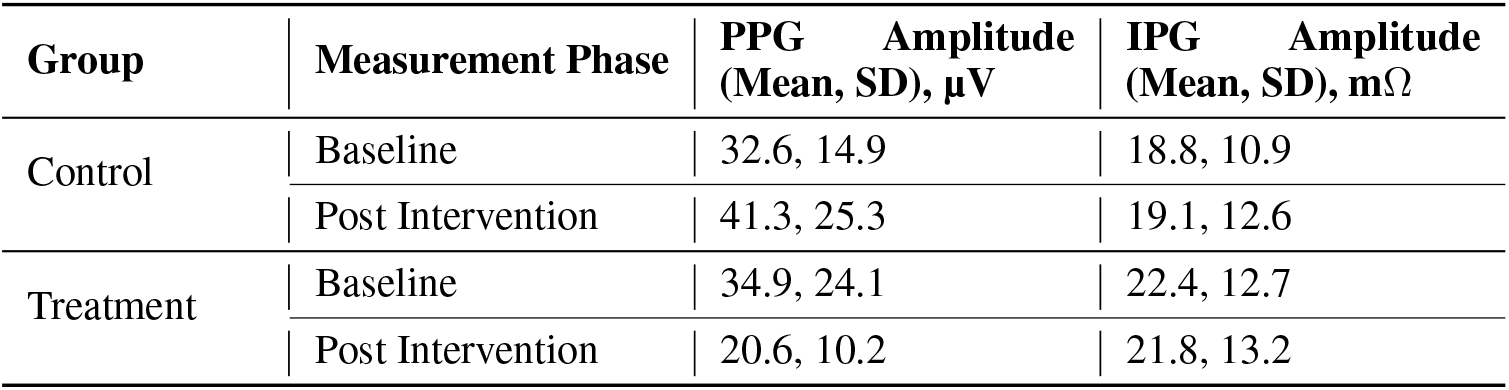
Aggregated signal amplitude analysis of all participants across respective groups and measurement phases.

| Group | Measurement Phase | PPG Amplitude<br>(Mean, SD), $\mu\text{V}$ | IPG Amplitude<br>(Mean, SD), $\text{m}\Omega$ |
| --- | --- | --- | --- |
| Control | Baseline | 32.6, 14.9 | 18.8, 10.9 |
|  | Post Intervention | 41.3, 25.3 | 19.1, 12.6 |
| Treatment | Baseline | 34.9, 24.1 | 22.4, 12.7 |
|  | Post Intervention | 20.6, 10.2 | 21.8, 13.2 |

**Table 3:** Paired t-test results for sensor amplitude metrics. Statistically-significant differences are indicated as follows: ns (not significant) for *p >* 0.05, * for *p <* 0.05, ** for *p <* 0.01, and *** for *p <* 0.001.

| Sensor | Metric | Group | Test type | t-stats | p-value | Stat. Sig. |
| --- | --- | --- | --- | --- | --- | --- |
| IPG_Z | Amplitude | control | baseline vs. post intervention | -0.29 | 0.77 | ns |
| IPG_Z | Amplitude | treatment | baseline vs. post intervention | 0.75 | 0.46 | ns |
| IPG_Z | $\Delta\text{Amplitude}$ | n/a | control vs. treatment | 0.61 | 0.55 | ns |
| PPG | Amplitude | control | baseline vs. post intervention | -3.1 | 0.006 | ** |
| PPG | Amplitude | treatment | baseline vs. post intervention | 3.6 | 0.002 | ** |
| PPG | $\Delta\text{Amplitude}$ | n/a | control vs. treatment | 3.8 | 0.001 | *** |

**Table 4:** Paired t-test results for skin temperature. Statistically-significant differences are indicated as follows: ns (not significant) for p > 0.05, * for p < 0.05, ** for p < 0.01, and *** for p < 0.001.

| Sensor | Metric | Group | Test Type | t-stats | p-value | Stat. Sig. |
| --- | --- | --- | --- | --- | --- | --- |
| IR thermal camera | Temp<sub>proximal</sub> | control | baseline vs. post intervention | -2.7 | 0.01 | * |
| IR thermal camera | Temp<sub>proximal</sub> | treatment | baseline vs. post intervention | 12 | 0.00 | *** |
| IR thermal camera | ΔTemp<sub>proximal</sub> | n/a | control vs. treatment | 12 | 0.00 | *** |
| IR thermal camera | Temp<sub>central</sub> | control | baseline vs. post intervention | -3.4 | 0.003 | ** |
| IR thermal camera | Temp<sub>central</sub> | treatment | baseline vs. post intervention | 14.2 | 0.00 | *** |
| IR thermal camera | ΔTemp<sub>central</sub> | n/a | control vs. treatment | 14 | 0.00 | *** |
| IR thermal camera | Temp<sub>distal</sub> | control | baseline vs. post intervention | -2.4 | 0.03 | * |
| IR thermal camera | Temp<sub>distal</sub> | treatment | baseline vs. post intervention | 19 | 0.00 | *** |
| IR thermal camera | ΔTemp<sub>distal</sub> | n/a | control vs. treatment | 18 | 0.00 | *** |

**Table 5:** Paired t-test results for blood pressure and heart rate. Statistically-significant differences are indicated as follows: ns (not significant) for *p >* 0.05, * for *p <* 0.05, ** for *p <* 0.01, and *** for *p <* 0.001.

| Sensor | Metric | Test Type | t-stats | p-value | Stat. Sig. |
| --- | --- | --- | --- | --- | --- |
| Arm cuff | SBP | start vs. end | 0.76 | 0.46 | ns |
| Arm cuff | DBP | start vs. end | 0.0 | 1.0 | ns |
| Arm cuff | HR | start vs. end | 2.3 | 0.03 | * |

A paired t-test analysis revealed a statistically significant difference (p = 0.001) in PPG signal amplitude between the treatment and control groups (Fig. 8, Table. 3). The treatment group exhibited a mean decrease of 14.3 µV in PPG amplitude (i.e. -41%), whereas the control group showed a mean increase of 8.7 µV (i.e. +27%). No statistically significant differences were observed in the change in IPG amplitude between the two groups (Fig. 7, Table. 3).

**Figure 7.**
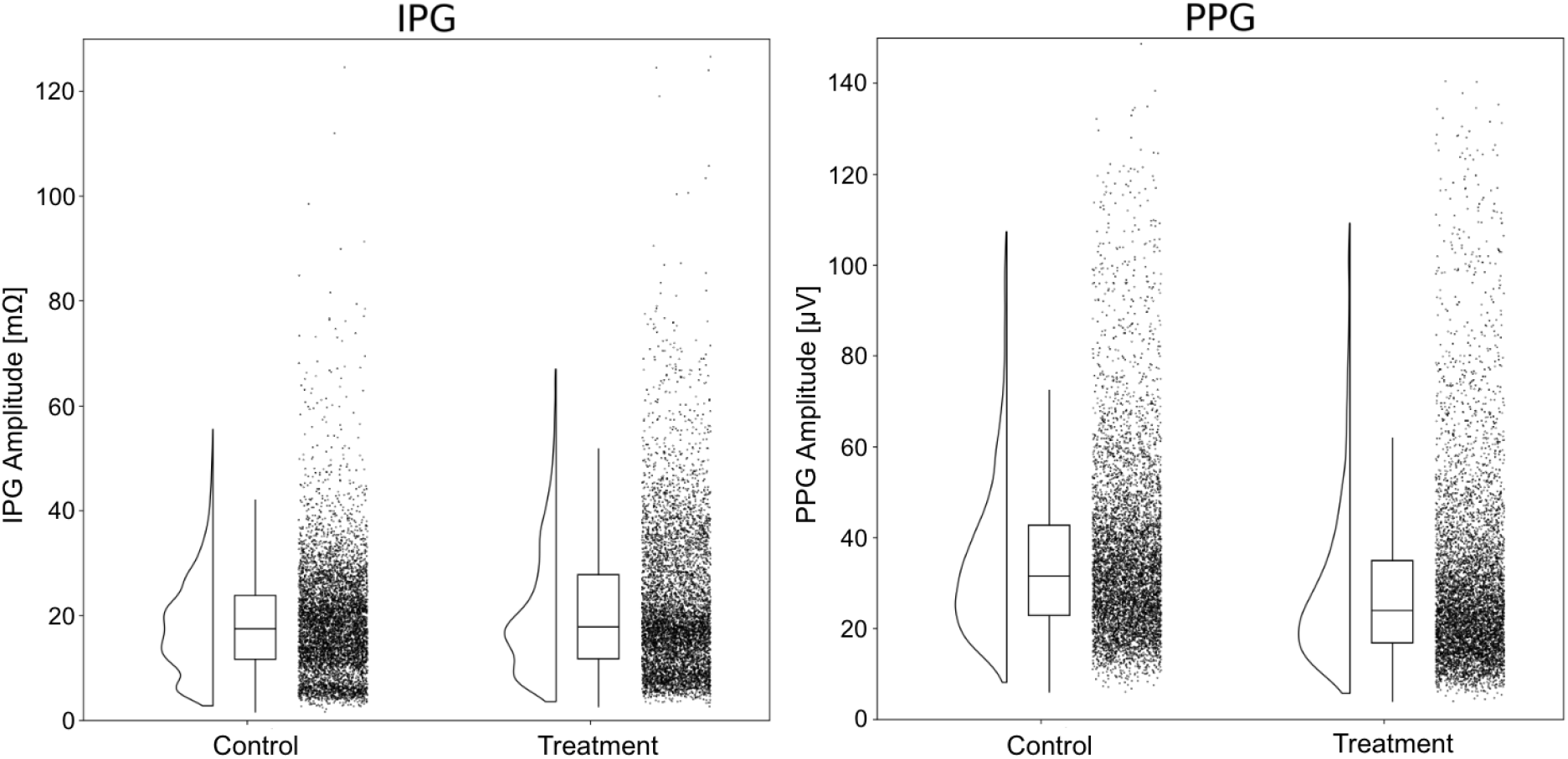
Distributions of the amplitudes from IPG (left) and PPG (right) sensors. Distributions represent all beats from the 12 min-long protocol for the control group and the treatment group respectively.

**Figure 8.**
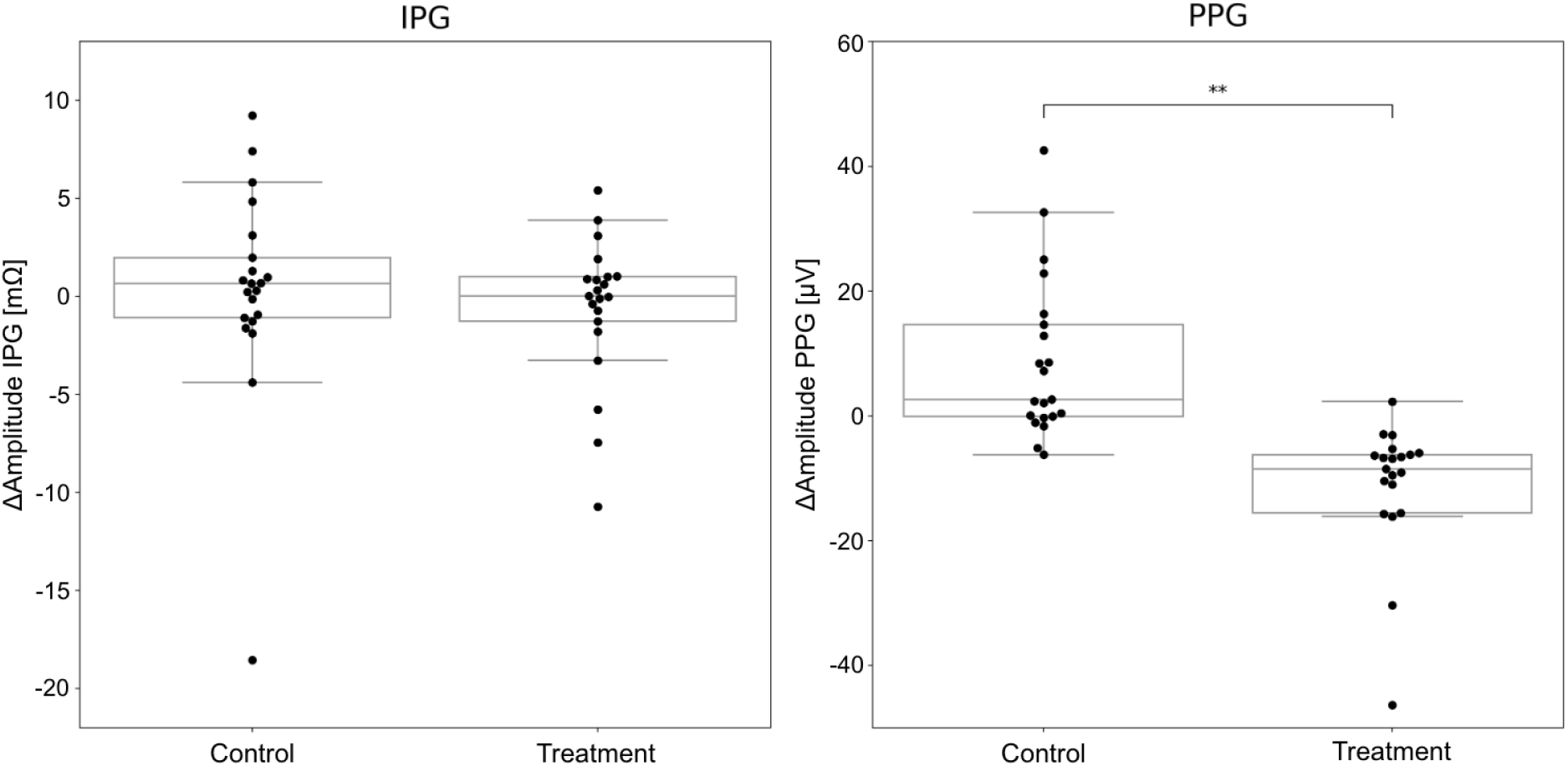
Changes in IPG and PPG amplitudes (post intervention - baseline) across control and treatment groups. ^**^ marks a statistically significant change of p < 0.01.

### 3.4 Timing Analysis

The timing analysis used the same peak finding approach described in 3.3 Signal Amplitude Analysis. Pulse arrival time (PAT) was also calculated for IPG and PPG respectively by finding the difference in time between the ECG R-peak and the valley/foot for PPG, and the valley/foot for IPG Z. The time difference between the valley of the IPG Z and the valley of PPG was calculated as well (Δ*T*_*xPG*_) [9]. Heart rate had a modest correlation (Pearson correlation coefficient, R = -0.55) with PPG-derived PAT (Fig. 9). No significant correlations were observed between BP and PAT. No statistically-significant differences in timing metrics were observed in a paired t-test analysis between groups (control and treatment), or with test type (baseline vs. post intervention) (Table 6, Appendix). As shown in Fig. 10, Δ*T*_*xPG*_ had modest correlations with SBP (R = -0.40) and heart rate (R = -0.31). Pulse wave velocity (PWV) was also evaluated by dividing subject’s heights with PAT, wherein the PPG-derived PAT based PWV and SBP had a correlation coefficient of 0.41. For other PAT converted PWVs, there wasn’t a meaningful change in correlation coefficients.

**Table 6:**
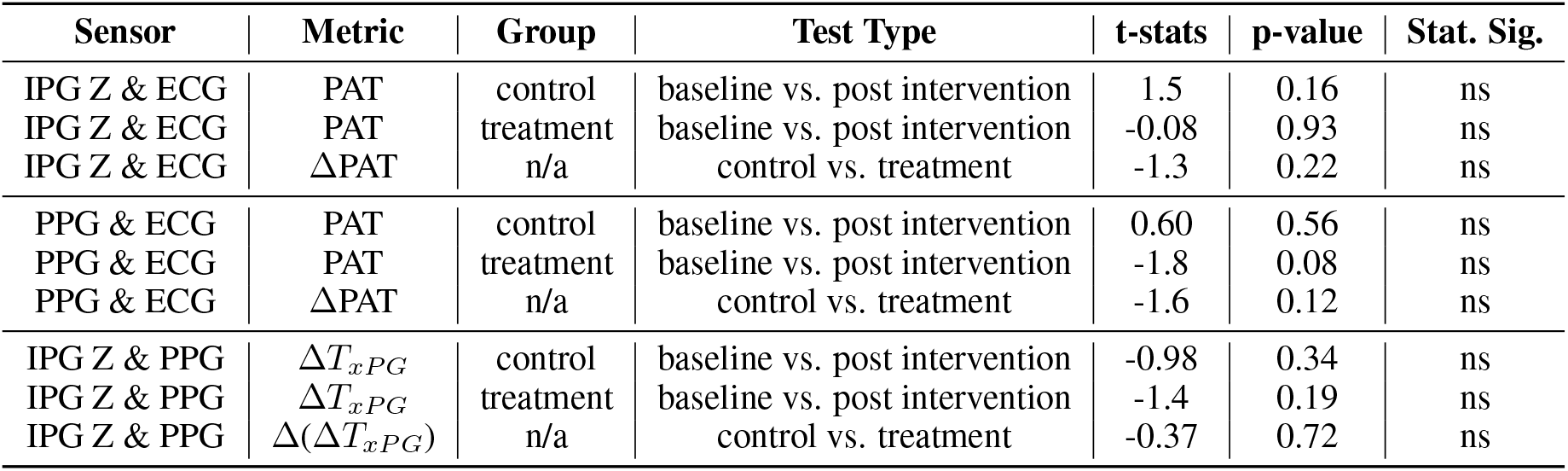
Paired t-test results for sensor timing metrics. Statistically-significant differences are indicated as follows: ns (not significant) for *p >* 0.05, * for *p <* 0.05, ** for *p <* 0.01, and *** for *p <* 0.001.

**Figure 9.**
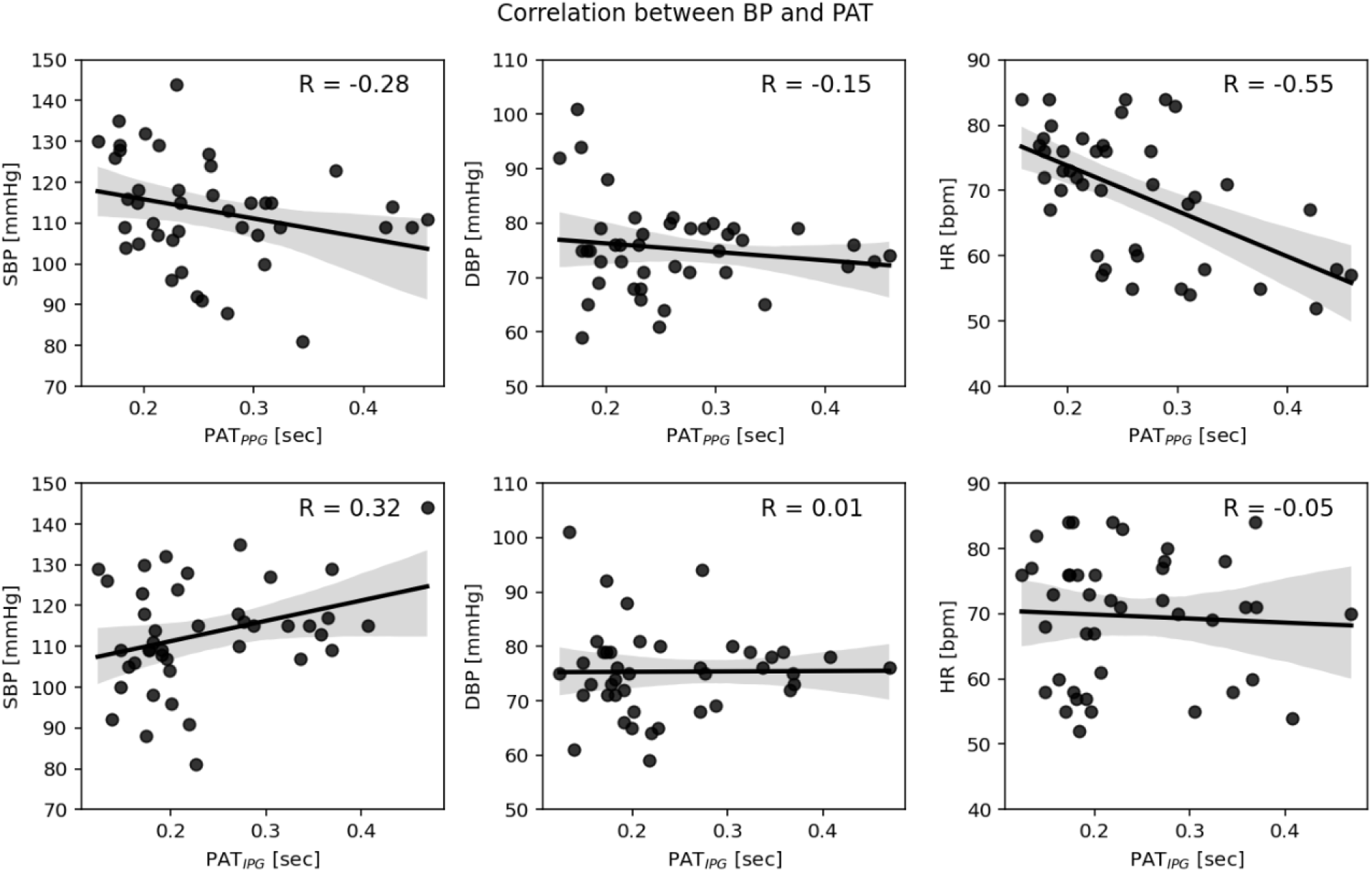
Correlation between Omron arm cuff measurements (SBP: left, DBP: middle, HR: right) and PAT (PPG-based: top, IPG-based: bottom). Pearson correlation coefficients (R) are marked.

**Figure 10.**
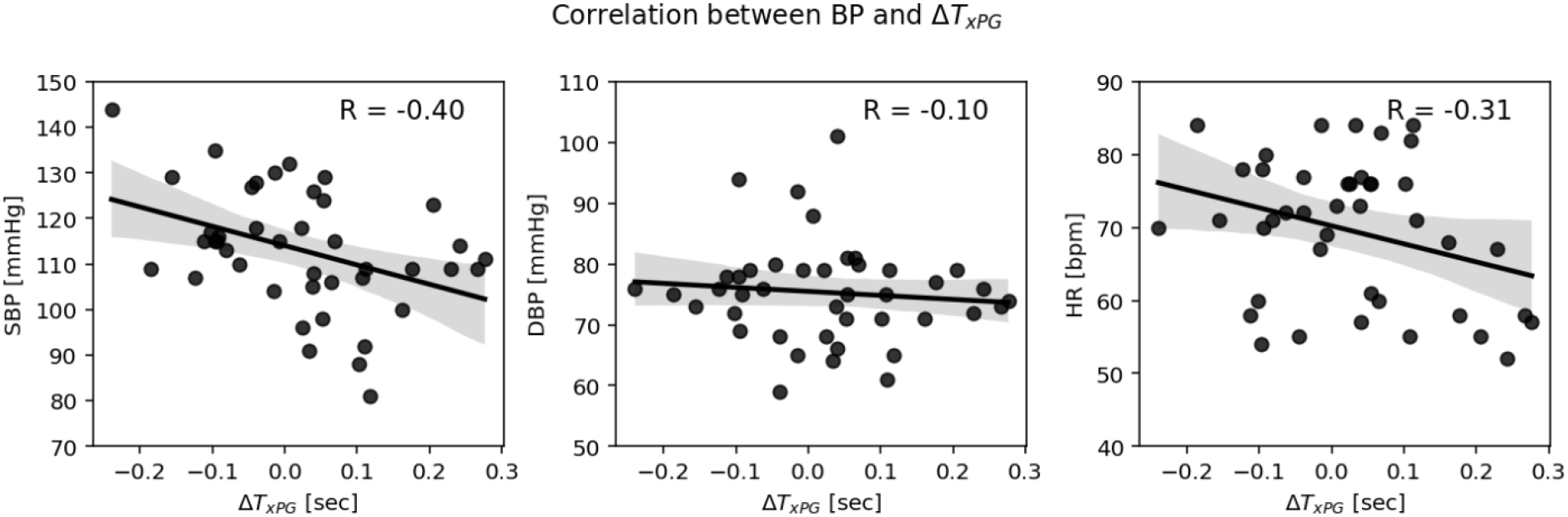
Correlation between Omron arm cuff measurements (SBP: left, DBP: middle, HR: right) and Δ*T*_*xPG*_. Pearson correlation coefficients (R) are marked.

## 4 Discussion

This study investigated the differential responses of IPG and PPG to temperature-mediated vasoconstriction, providing valuable insights into their distinct sensitivities to superficial blood flow dynamics. The within-subject experimental design, incorporating a controlled localized cooling intervention, enabled a direct and reliable comparison between the two sensing modalities. Using both real and fake ice cubes in the treatment and control groups respectively, it helped isolate the impact of temperature variation whilst mitigating potential confounding factors related to physical sensor disturbance.

As hypothesized, the PPG signal exhibited a statistically significant reduction in amplitude following localized cooling. This observation aligns with the established literature that highlights PPG’s responsiveness to microvascular alterations [6]. The increase in PPG amplitude during the control procedural group could be due to vasodilation coming from localized heating [8]. Conversely, IPG signals exhibited no statistically significant change in amplitude despite the induced vasoconstriction. This finding suggests that IPG is less sensitive to capillary-level changes and may primarily reflect blood flow in larger and/or deeper vessels.

The observed stability of IPG amplitude during vasoconstriction necessitates a critical examination of the underlying physiological mechanisms, particularly regarding the depth of signal penetration. There were commonalities in sensor placement and the cold intervention. Both PPG and IPG sensors were placed on top of the radial artery. The sensor locations (proximal, central, and distal) all showed a statistically significant decrease in skin temperature meaning the temperature drop was homogenous across the sensing sites.

Our hypothesis that IPG is primarily sensing a deeper blood flow aligns with the observation from the simulation based measurements of radial artery bioimpedance [10]. With the 3D model of human forearm fragment created in Comsol Multiphysics and inline Ag/AgCl electrodes placed on a radial artery similar to our setup, the finite element method simulation showed that part of the injected current from the outer electrodes directly flows through the radial artery, wherein the inner electrodes sense the voltage potential between them. The human forearm model used in the simulation was created using MRI-scan-based geometry and known skin, fat, bone, muscle, and tissue conductivity distributions.

Similarly, our hypothesis on the PPG primarily sensing a shallow blood flow aligns with the observations from prior works of high-speed imaging PPG [4], [11]. A common belief that PPG directly senses pulsating arteries is debunked. The PPG waveform rather reflects elastic deformations of the capillary bed. With the pulse oscillations of the arterial transmural pressure, the connective-tissue components deform resulting in periodic changes of both the light scattering and absorption. In other words, PPG indirectly monitors arterial pulsations by the sensing deformation of the shallow capillary beds.

No significant variations in PAT were observed for either IPG or PPG following localized cooling. PAT is sometimes used as a proxy for arterial stiffness and blood pressure fluctuations [12]. The absence of significant PAT changes suggests that the localized cooling may not have induced substantial changes in arterial stiffness or systematic blood pressure within the observed timeframe, which is consistent with the stable blood pressure measurements obtained via oscillometry. However, a statistically-significant decrease in heart rate was observed, likely due to participants sitting still for a relatively long period of time and asking them to relax. These findings suggest that the localized vasoconstriction primarily affected the microvasculature, with minimal impact on larger arterial compliance and overall systemic hemodynamics.

There are several limitations to this study. Firstly, the participant cohort consists primarily of Google employees located in the San Francisco Bay Area, potentially limiting the generalization of the findings to the broader population. Future studies should aim for greater demographic diversity. Secondly, further investigations into the signal penetration of PPG and IPG in these interventional scenarios with smaller form factors seems a natural extension of this work. Additionally, PPG and IPG sensors could be further optimized to better accommodate wearable form factors. For instance, multipath PPG could be considered instead of single channel PPG. Additionally, electrode design parameters for IPG might also be further optimized such as its positioning and product-friendly material choice.

## 5 Conclusion

This study investigated the impact of reduced capillary blood flow on IPG and PPG signals. The findings suggest that IPG signals, unlike PPG signals, remain relatively stable despite significant changes in capillary blood flow due to cold-induced vasoconstriction. This observation has implications for the potential use of IPG in wearable devices for monitoring deeper blood flow and cardiovascular activity wherein IPG can provide a valuable complementary measure to PPG. This information can be used to develop more robust and accurate models related to blood flow, paving the way for more personalized and adaptive human-computer interactions.

## Credit Authorship Contribution Statement

Seobin Jung, Seamus Thomson: Writing – original draft, Visualization, Software, Methodology, Investigation, Formal analysis, Data curation, Conceptualization. Alexandros Pantelopoulos: Writing – review and editing, Supervision, Software, Conceptualization. Lindsey Sunden: Supervision, Conceptualization. Pete Richards: Writing – review and editing, Supervision. Shwetak Patel, Sam Sheng: Review, Funding acquisition.

## Acknowledgement

The authors would like to thank Brent Winslow and Tracy Giest for their help with preparing the IRB application. The authors also appreciate Conor Heneghan for his help reviewing this paper.

## Data Availability

The datasets generated during and/or analysed during the current study are available from the corresponding authors on reasonable request.

## A Appendix

